# Semi-supervised identification of SARS-CoV-2 molecular targets

**DOI:** 10.1101/2021.05.03.440524

**Authors:** Kristen L. Beck, Ed Seabolt, Akshay Agarwal, Gowri Nayar, Simone Bianco, Harsha Krishnareddy, Vandana Mukherjee, James H. Kaufman

## Abstract

SARS-CoV-2 genomic sequencing efforts have scaled dramatically to address the current global pandemic and aid public health. In this work, we analyzed a corpus of 66,000 SARS-CoV-2 genome sequences. We developed a novel semi-supervised pipeline for automated gene, protein, and functional domain annotation of SARS-CoV-2 genomes that differentiates itself by not relying on use of a single reference genome and by overcoming atypical genome traits. Using this method, we identified the comprehensive set of known proteins with 98.5% set membership accuracy and 99.1% accuracy in length prediction compared to proteome references including Replicase polyprotein 1ab (with its transcriptional slippage site). Compared to other published tools such as Prokka (base) and VAPiD, we yielded an 6.4- and 1.8-fold increase in protein annotations. Our method generated 13,000,000 molecular target sequences— some conserved across time and geography while others represent emerging variants. We observed 3,362 non-redundant sequences per protein on average within this corpus and describe key D614G and N501Y variants spatiotemporally. For spike glycoprotein domains, we achieved greater than 97.9% sequence identity to references and characterized Receptor Binding Domain variants. Here, we comprehensively present the molecular targets to refine biomedical interventions for SARS-CoV-2 with a scalable high-accuracy method to analyze newly sequenced infections.

## 1 Introduction

The ongoing SARS-CoV-2 pandemic has undoubtedly shaped our lives as one of the most significant global health challenges we are facing. However, unlike previous pandemics, we now have sequencing technology with tremendous throughput to analyze the genomic content of SARS-CoV-2. As labs around the world sequence isolates from infected individuals, we can track and characterize the viral genome evolution in almost near real-time as the pandemic keeps infecting the worldwide population.

The first sequenced SARS-CoV-2 genome (*1*) was submitted to NCBI January 17, 2020 and has become the accepted reference standard commonly referred to as the Wuhan reference genome (NCBI RefSeq ID: NC 045512.2). Since that point, tens of thousands of genomes are published on a weekly basis. The SARS-CoV-2 genome is comprised of a 29,000 base pairs (bp) single-stranded RNA (38% GC content) with four structural proteins, two large polyproteins which are cleaved to form non-structural proteins, and several accessory proteins (*2, 3*). There are two overlapping open reading frames responsible for Replicase polyprotein 1a (pp1a) and Replicase polyprotein 1ab (pp1ab) which yield the longest products from the genome and the majority of the non-structural proteins.

In comparison to other coronaviruses, SARS-CoV-2 differs phenotypically with its significant increase in transmissibility and asymptomatic or presymptomatic transmission, as well as genotypically from its polybasic cleavage site insertion in the S protein (*4*). However, it maintains several other *Coronaviridae* traits such as gene order consistency and transcriptional slippage (*5*). The -1 programmed ribosomal frameshift responsible for transcriptional slippage has been observed to occur at the point where ORF1 (responsible for pp1a) continues as ORF2 (responsible for pp1ab) and is defined by an RNA signature marking the slippery site (*6*). This phenomenon allows the virus to control the relative levels of its protein expression (*6*) and may be useful in therapeutic targeting to limit protein production. Additionally, this unique trait creates a challenge for traditional bioinformatic genome annotation programs which assume that the more typical continuous 5’ to 3’ translation can be effectively used to form the correct protein sequence which is not the case for SARS-CoV-2.

There are several viral genome annotation methods such as VAPiD, Prokka, InterProScan, and others (*7–9*) that aim to provide autonomous (that is, no reference genome required) annotation of genes and proteins. Some of these tools have issued special releases to aid in annotating SARS-CoV-2 genomes. Yet, many of these tools do not provide sufficient accuracy with “off the shelf” use and have not yet been applied at scale as the available SARS-CoV-2 sequence data grows. Additionally, several variants of SARS-CoV-2 genomes have emerged including the D614G (*10*) variant, which appeared earlier in the pandemic, or the more recent B.1.1.7 variant (*11, 12*), which now represents the majority of new cases in the USA (*13*). The mutations defining these variants can present challenges for complete automation of genome annotation and this can be further exacerbated by the SARS-CoV-2 transcriptional slippage site.

As an alternative to an autonomous genome annotation method, alignments to the Wuhan reference genome (*1*) using tools such as NextStrain’s Augur (*14*), Bowtie2 (*15*), or UCSC SARS-CoV-2 genome browser (*3*) can be completed. This type of supervised analysis uses published gene coordinates to extract sequences from the query genome based on positional and sequence similarity to a reference genome. However, this creates a considerable dependency on a single reference genome. Since it is currently estimated that SARS-CoV-2 typically mutates approximately twice per month on any given transmission chain (*16*) and can be subject to recombination events (*17*), a reference-guided approach may face limitations as the virus continues to evolve or increases the rate at which it evolves.

In this work, we present a semi-supervised custom pipeline to annotate all genes, proteins, and functional domains for SARS-CoV-2. This method has been applied to 66,905 SARS-CoV-2 genomes collected from NCBI GenBank (*18*) and GISAID (*19*). This approach yielded nearly 13 million new molecular sequences and connections that can be accessed through the IBM Functional Genomics Platform, a tool made freely available to the global research community (*20*). With our method, we achieved 98.5% average protein set membership identification accuracy and an average observed over expected protein length ratio of 99.1%. Additionally, in comparison to other tools such as Prokka or VAPiD, we identified 6.4- and 1.8-fold more protein annotations, respectively. Furthermore in a targeted analysis, we achieved greater than 97.9% sequence identity in spike glycoprotein domains.

To illustrate the value of this approach, we utilized the variants identified in this collection to track the emergence of the D614G and N501Y spike glycoprotein variants over time and by region of exposure. The complete collection of SARS-CoV-2 genes, proteins, and functional domains continues to be updated and can be accessed via a web browser user interface or our developer toolkit.^1^ Ultimately, we present a comprehensive comparative analysis and data resource of 66,905 publicly available SARS-CoV-2 viral sequences, with the aim of identifying potential targets to aid in vaccine, diagnostic, or therapeutic development.

## 2 Results

Here, we present a novel semi-supervised pipeline to annotate gene, protein, and functional domain molecular targets from SARS-CoV-2 genomes and demonstrate the resulting accuracy against known reference data and other bioinformatic tools (see Methods). This pipeline provides improvements over base Prokka and InterProScan by adding a novel capability that more accurately processes sequences the slippery site junction in Replicase polyprotein 1ab to identify the correct sequence which is absent or artificially truncated in other methods. Additionally, we incorporated a targeted search for three key proteins: ORF9b, ORF10, and Envelope small membrane protein which would otherwise be missing in the protein annotation set. We show that our pipeline has an improvement on annotation accuracy by 1.8- and 6.4-fold compared to VAPiD and base Prokka. We also evaluated genome quality from data in public repositories and quantitatively evaluated commonly used quality criteria for their effects on the resulting annotations.

### 2.1 Assessment of SARS-CoV-2 genome quality in multiple data sources

For effective genome annotation, an important first step is to assess the quality of the input genomes. In this study, we analyzed a corpus of 66,905 SARS-CoV-2 genomes (Supplementary File 1) deposited over the span of eight months from 108 countries into two key aggregate data sources: GISAID (*19*) and NCBI GenBank (*18*). We observed an average of 0.0067% unknown bases (denoted as N per IUPAC definitions) per genome (range 0–46.76%), and all genomes were observed to have less than 1% degenerate bases (Figure 1a). The presence of unknown bases can indicate insufficient genome coverage or other issues from genome assembly. Next, we aimed to identify criteria for inclusion of a SARS-CoV-2 genome assembly into our platform to ensure that input data for molecular target identification is of the highest quality. Therefore, we evaluated two commonly used criteria for their effects on prediction of full length protein sequences. Briefly, Criteria A is more permissive and prioritizes the ratio of length vs. coverage whereas Criteria B is more stringent and applies a higher penalty to the number of gaps (detailed definitions in Section 4.1). Here, “full length” is defined as a protein sequence length within 10% of the known UniProt protein reference sequence indicated in the SARS-CoV-2 proteome as defined in ViralZone (*21, 22*). Figure 1b demonstrates that Criteria A yielded the highest count of full length protein products (35,099 non-redundant protein sequences) while also effectively reducing the majority of truncated products (2,197 non-redundant protein sequences). If applying the more stringent Criteria B, over 14,000 high quality protein products would be inadvertently removed. Based on this, we proceeded with applying the thresholds defined by Criteria A to the corpus of genomes analyzed in this work. From GISAID and GenBank genomes, 9.9% (out of 55,708) and 3.1% (out of 11,197) of genomes fell below this criteria (Figure 1a) and are removed from subsequent analysis and marked as inactive unless otherwise mentioned.

**Figure 1:**
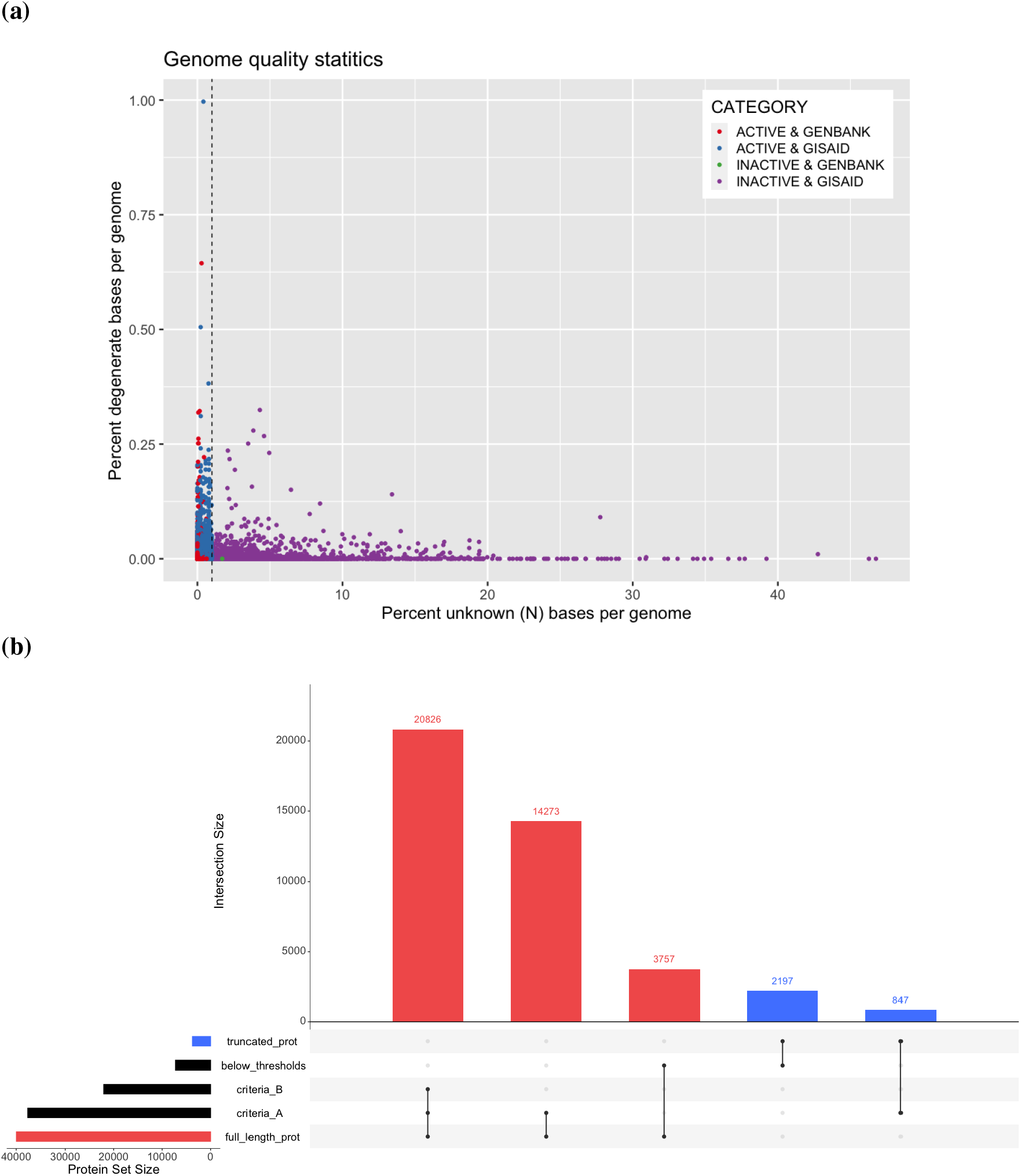
SARS-CoV-2 genome quality observations and their effect on annotation results. The percent of unknown (N) and degenerate bases (as defined by IUPAC) are calculated as a function of total genome size for SARS-COV-2 genomes from two sources: NCBI GenBank and GISAID (**1a**). The quality threshold of unknown bases is indicated with a dashed line. Also, full length (red) or truncated (blue) protein products are indicated for genomes by quality criteria status: our selected criteria (Criteria A), a more stringent criteria (Criteria B), or below quality thresholds (**1b**). For criteria definitions, see Section 4.1.

As part of our other pre-processing steps for genomic data (*20*), we computed md5 hashes of all genome sequences to track duplicates. The rate of ‘duplicated’ SARS-CoV-2 genome sequences within a single data source or between data sources is indicated in Table 1. From the data at hand, it is unclear whether these duplicated genome sequences as others have observed (*23–25*) are artefacts of data processing e.g. due to alignments to a single reference genome or are a result of sampling multiple patient infections within the same lineage. Of the 10,528 genomes that are duplicated within GISAID, we compared the metadata for each entry: 3,953 are described with matching metadata entries in addition to identical full length genome sequences and therefore may be more likely to be data duplication artefacts. These potential duplication events are reported here but are not removed from subsequent analysis.

**Table 1:** Distribution of Duplicated Genome Sequences. The number of unique genome sequences (count of non-redundant md5 hashes) and total genome accessions (count of all entries) are listed for each source: GISAID, GenBank, and NCBI RefSeq. Then pairwise comparisons are made for each genome sequence to indicate how many times a genome sequence is duplicated (exact md5 match of genome sequence) within its source or between sources and for how many entries this accounts for (count of genome accessions). For RefSeq comparisons, note only one SARS-CoV-2 reference genome sequence is available NC 045512.2

| Source 1 | Source 2 | Genome Sequence Hashes | Genome Accessions |
| --- | --- | --- | --- |
| GISAID | NA | 47,908 | 55,708 |
| GENBANK | NA | 9,398 | 11,196 |
| REFSEQ | NA | 1 | 1 |
| GISAID | GISAID | 2,791 | 10,528 |
| GISAID | GENBANK | 5,559 | 13,977 |
| GENBANK | GENBANK | 706 | 2,504 |
| GISAID | REFSEQ | 1 | 43 |
| GENBANK | REFSEQ | 1 | 11 |

### 2.2 Quantification of protein sequence prediction accuracy

For an autonomous COVID-19 genome annotation pipeline to achieve clinical and biological relevance, it must accurately identify all known molecular targets within a genome. The SARS-CoV-2 proteome (*2,22*) is defined as having fourteen protein products each with a corresponding gene sequence present in each genome. SARS-CoV-2 proteins are split into structural and non-structural, but all proteins are required for the virus to carry out its life cycle which includes host cell invasion, replication, and transmission (*26*). Using our gene and protein annotation method (Section 4.2), we achieved an average per protein identification accuracy of 98.5% ± 2.9% across all genomes above the aforementioned quality thresholds. The number of observations per protein (Figure 2a) indicates that we are able to achieve complete or near-complete protein set membership for all genomes. Each protein is a translated gene sequence, and thus the equivalent gene identification accuracy is also achieved.

**Figure 2:**
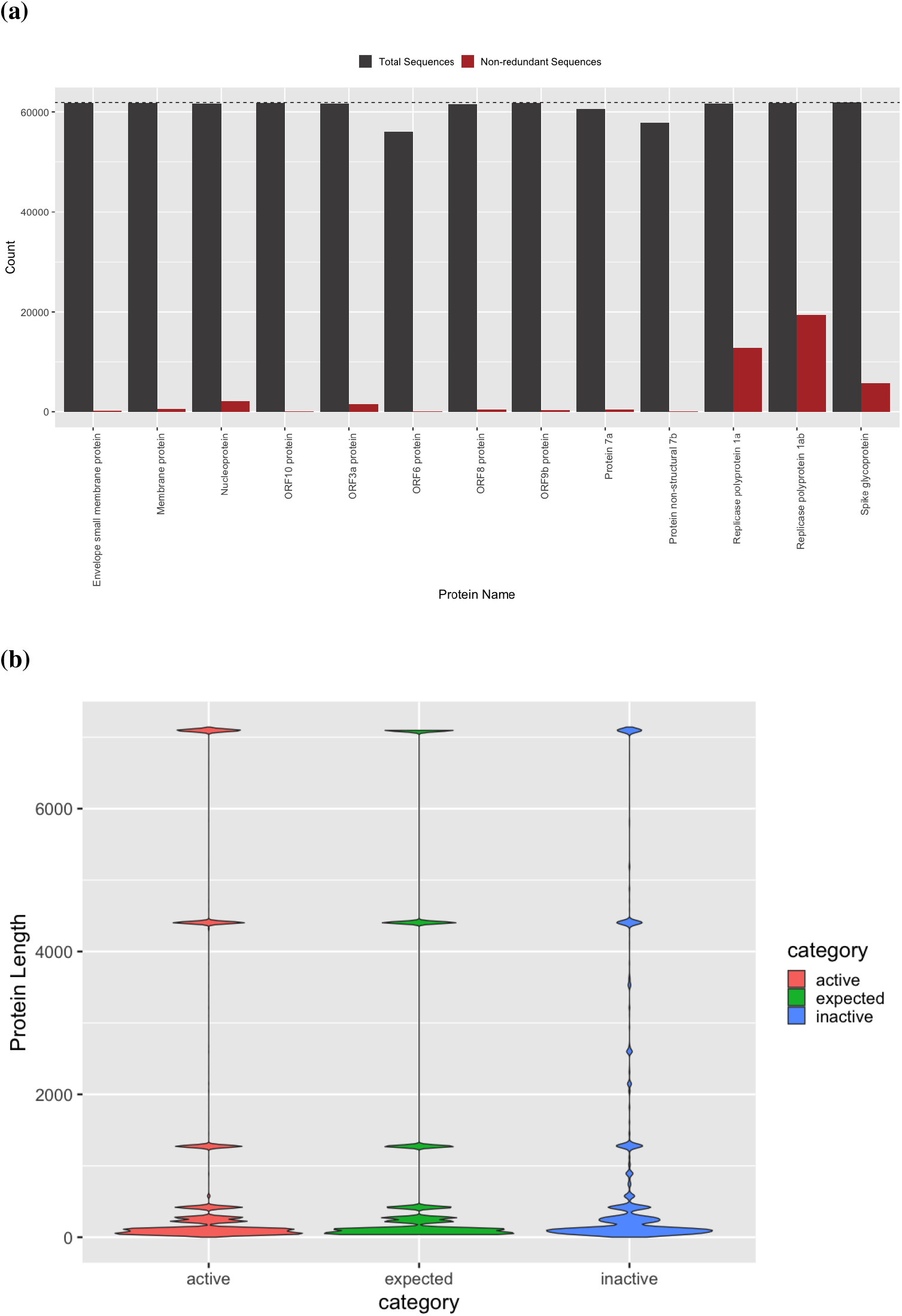
Protein annotation set membership and sequence accuracy. Set membership of all (grey) and distinct (red) protein sequences identified in SARS-CoV-2 genomes above quality thresholds (**2a**). Count indicates the number of times a given protein was observed in the entire active genome corpus (dashed line). (**2b**) Protein size distribution of known protein references (expected, green) more closely matches our *in silico* predicted protein sequences from genomes above quality thresholds (active genomes, red) compared to products predicted from genomes below quality thresholds (inactive genomes, blue).

Furthermore, not only must the complete set of named genes and proteins be identified for accurate genome annotation, but the generated sequences must also be grounded in biological reality. Specifically, *in silico* predicted sequences should not be truncated with respect to the length of known references, and the mutational density must be low considering the temporally recent emergence of SARS-CoV-2 and observed lower mutation rate compared to other RNA viruses (*16*). Using our semi-supervised gene and protein annotation method (Section 4.2), we were able to identify full length protein products that on a per protein basis match expected lengths of known reference sequences with an average observed / expected protein length value of 99.1% (Figure 2b). The distributions of our predicted and the expected protein sequence lengths are observed to be statistically similar by two-sample Kolmogorov–Smirnov test (D = 0.0071, p *<* 2.2e-16) and are 8.75-fold more similar (D = 0.0617) than those predicted from genomes not passing our quality thresholds, i.e., inactive genomes.

In addition, certain gene and protein sequences required us to develop additional targeted methodological advances for identification (Section 4.2.2). Specifically, Replicase polyprotein 1ab (pp1ab) is the longest gene sequence within SARS-CoV-2 and its protein sequence is cleaved into 16 non-structural proteins (*2*). It overlaps with Replicase polyprotein 1a, and during translation, undergoes -1 programmed ribosomal frame shift at what is known as a slippery site (*6*). Both of these attributes make it more challenging to accurately identify with off-the-shelf *in silico* genome annotation methods. Therefore, we implemented a semi-supervised method (Section 4.2.2) to correct and extend the putative predicted gene coordinates for pp1ab and adjust the translation method to accommodate ribosomal frame shift which is a problem that negatively affects other bioinformatic tools. Our algorithmic improvement yielded full length pp1ab sequences in all genomes with greater than 95% sequence identity to the reference pp1ab sequence (UniProt ID: P0DTD1) in over 99.15% of the variants we predict (Figure 3).

**Figure 3:**
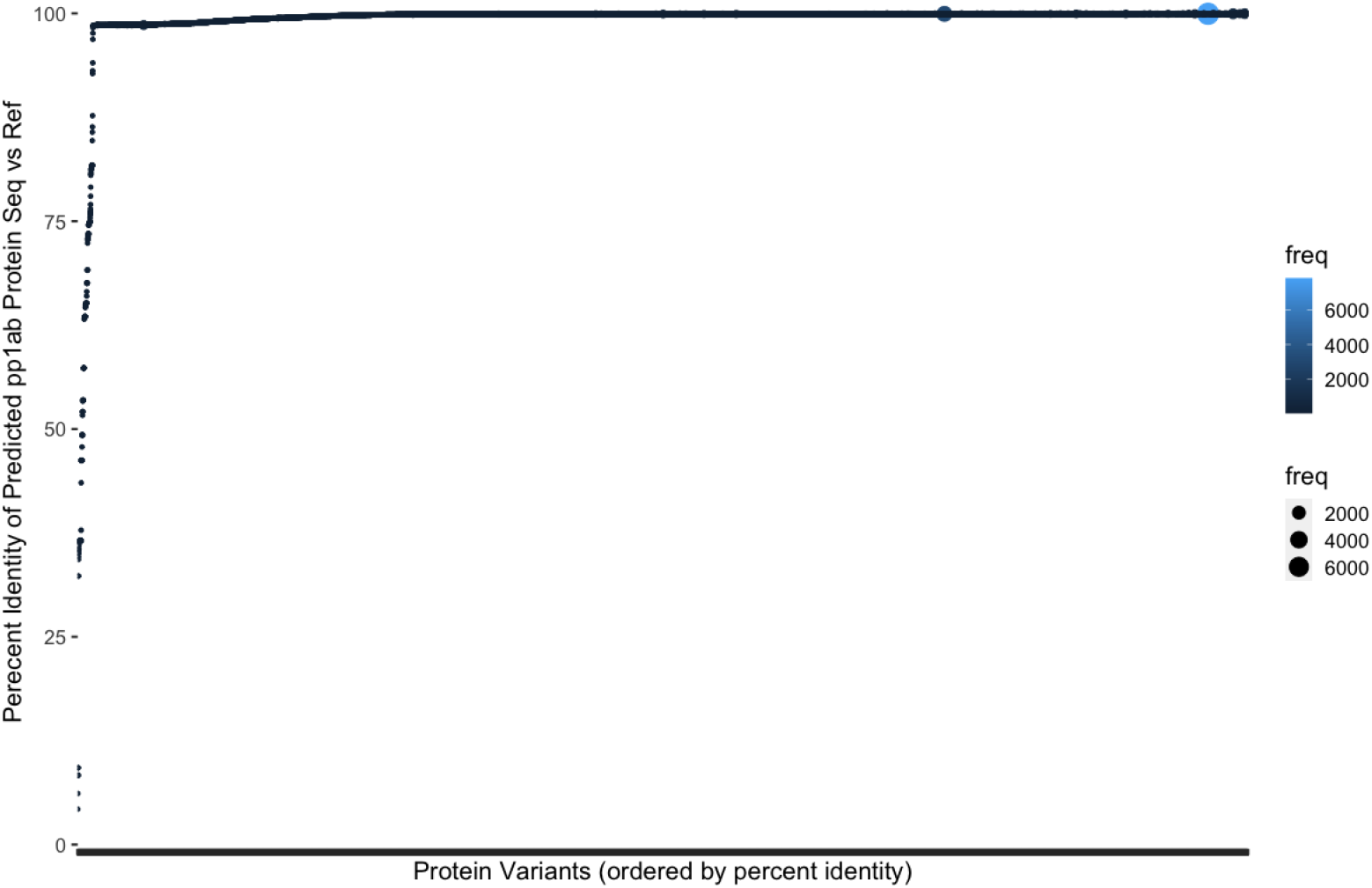
For Replicase polyprotein 1ab, the variant frequency and sequence similarity of our predicted protein to the known UniProt ID:P0DTD1 references is shown

Ultimately from this genome corpus, we were able to identify over 13M gene, protein, and functional domain sequences in total (Table 2). Our system only stores each uniquely identified sequence once (distinct sequence), but maintains the relationship to its originating genome and to any connected sequences e.g. gene, protein, or domain sequence providing the total sequences identified (Table 2).

**Table 2:** Observed gene, protein, and functional domain biological entities in total count (redundant) and unique sequences (distinct) in active genomes.

| Type | Total Count | Unique Count |
| --- | --- | --- |
| Gene | 936,603 | 59,531 |
| Protein | 815,878 | 42,611 |
| Domain | 11,621,784 | 59,271 |

### 2.3 Comparative analysis of genome annotation methods

With regard to pipeline accuracy, we benchmarked our pipeline against VAPiD (*7*), which has created a special release for annotating SARS-CoV-2 genomic data, and Prokka (*8*), a prokaryotic genome annotation tool for bacteria and virus. From the same set of genomes, we contrasted the resulting protein annotations (Figure 4) in the context of set membership as well as in observed protein sequence length compared to reference protein sequence length (expected). VAPiD and our method both achieved high accuracy with regard to truncated proteins, but our pipeline elicited more proteins in the highest accuracy category and 1.8-fold more protein annotations overall (Figure 4). Prokka, on the other hand, did not yield any full length pp1ab protein sequences and generated a high amount of missing or truncated proteins (Figure 4). Our method was able to identify 6.4-fold more protein products compared to base Prokka and was able to generate full-length pp1ab products with high sequence identity to known UniProt references.

**Figure 4:**
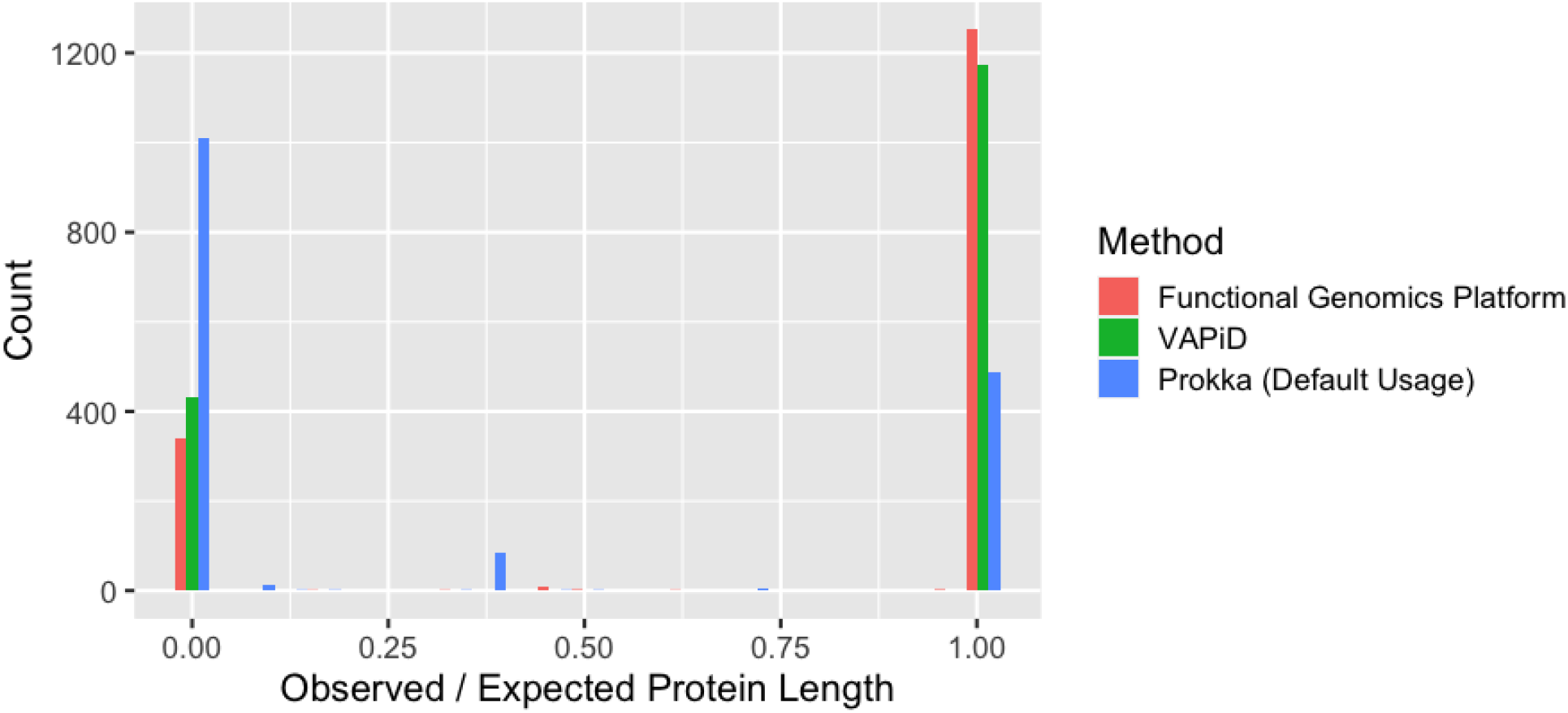
Protein length comparisons against known reference sequences for three pipelines: Functional Genomics Platform (our method), VAPiD, Prokka with default usage for viral genomes. The count of protein sequences at each observed / expected value is plotted for each pipeline. Length is set to zero if a protein is missing in the results from that pipeline but present in another.

### 2.4 Quantification of domain sequence prediction accuracy

To evaluate functional domain annotation accuracy, we analyzed functional domains for spike glycoprotein (S protein) as this is one of the most studied proteins in SARS-CoV-2 and it is of high biomedical importance. Specifically, we analyzed the set membership completeness of our predicted domains against the domain architecture indicated by InterProScan for the UniProt RefSeq P0DTC2 (https://www.ebi.ac.uk/interpro/protein/reviewed/P0DTC2/). From 5,702 distinct spike protein sequences, our annotation method yielded an average 94.4% ± 4.4% domain set membership accuracy (Figure 5a) with only two unexpected domain annotations (IPR043002 and IPR043614) found in less than 0.1% of the proteins. For IPR043473, the domain architecture is split across two locations which accurately accounts for the number of observations of this domain being greater than the number of spike proteins analyzed. The corresponding count of distinct sequences for each domain is indicated in Figure 5a and these domains were observed to have 679 unique sequences on average. The S1-subunit of the N-terminal domain for SARS-CoV (IPR044341) and Betacoronaviruses (IPR032500) were observed to have the highest count of non-synonymous variants. Furthermore, we calculated the amino acid percent identity of each of our predicted domain sequences against domain sequences extracted reference S protein, and all domains achieved greater than 98% median percent identity (Figure 5b). Together, this indicates completeness of annotation and correctness of the predicted domain sequences.

**Figure 5:**
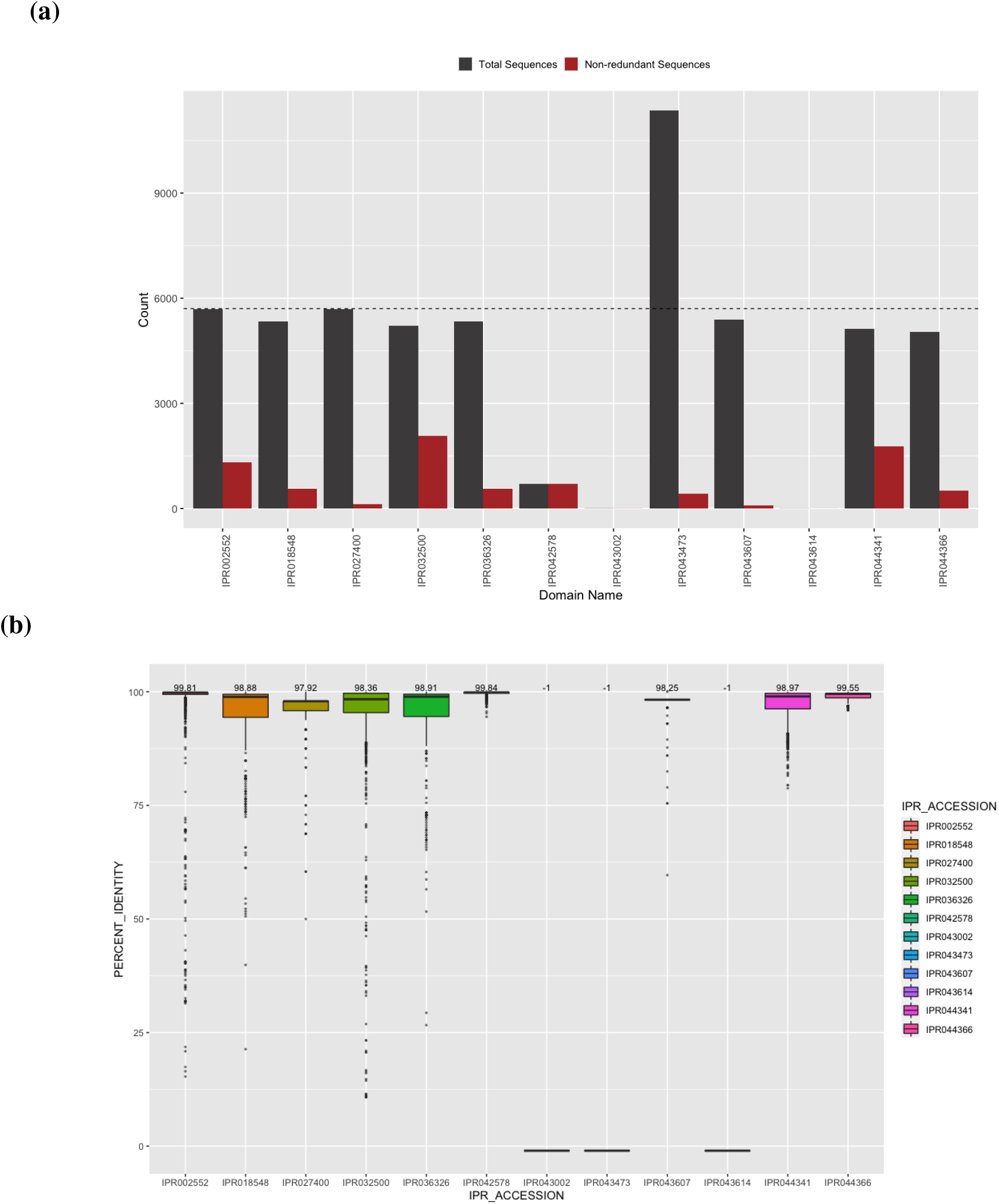
Domain annotation set membership and sequence accuracy for spike glycoprotein. Set membership of all (grey) and distinct (red) domain sequences identified in spike glycoprotein (**5a**). Count indicates the number of times a given domain was observed for all S proteins (dashed line). (**5b**) indicates the percent identity of our predicted domain sequences against reference domain sequences where possible to be calculated. In the absence of a reference sequence, percent identity is indicated as -1.

### 2.5 Distributions of variants shift over time

We identified the exhaustive set of genes, proteins, and functional domains (Table 2) for the corpus of genomes indicated previously. The number of variants (distinct sequences) differs per molecular target across all bio-entities as well each variants’ frequency (cumulatively shown in the redundant count). As SARS-CoV-2 undergoes mutation events, a comprehensive catalog of variants is essential for developing molecular interventions with sufficient specificity and binding efficiency. We observe a median of 425 unique sequences with non-synonymous mutational differences per protein (the number of unique sequences range=109–19,406) per protein. The S protein which is involved in invasion of human cells through interaction with ACE2 (*27*) is observed to have the highest number of variants among structural SARS-CoV-2 proteins (Figure 2a). Not surprisingly, the non-structural products of ORF1a and ORF1ab are also observed to have a higher amount of sequence variants compared to other SARS-CoV-2 proteins (Figure 2a).

Since the S protein is the key gatekeeper of host cell invasion and the target of multiple vaccines, antivirals, and diagnostics, we further examined its observed variants. We observed two predominant S protein variants that shift in their cumulative frequency over time (Figure 6a). Initially, an exact match to the reference spike glycoprotein sequence (green line, UniProt: P0DTC2) is observed most frequently. Then in mid-April, the notable variant D614G (orange line) with now known increased infectivity due to interaction with ACE2 receptor (*28, 29*) overtakes the ancestral reference sequence (green line) in its abundance achieving fixation. Two other differing sequences are observed at lower abundance in this genome cohort and correspond to P1140X (olive line) and S2 cleavage product (pink line). Additionally, there are minor variants observed in less than 1% of sequenced genomes. For example, we observe 5 protein sequences to contain the N501Y mutation from 13 genomes originating from Oceania (submitted 2020-07-02) and North America (submitted 2020-06-01). These variants are also observed to contain the D614G mutation, but are not present with the 69-70 deletion present in the B.1.1.7 variant of concern (UK) or B.1.351 E484K mutation (South Africa). Our observations are consistent with the current understanding of multiple introduction events causing the emergence of the N501Y variant (*30*). Furthermore, since this variant is observed in the B.1 and B.1.1 lineages which predate the current B.1.1.7 and B.1.351 variants it further clarifies the current timing of mutational introduction points in the pandemic. Some of our observed sequence variants may be due sampling to limitations or data artefacts e.g. sequencing or genome assembly error, but if a minor variant confers a selective advantage, its frequency could shift to become a more common variant as we have seen in recent months with the B.1.1.7, P.1, and B.1.351 variants (*31*).

**Figure 6:**
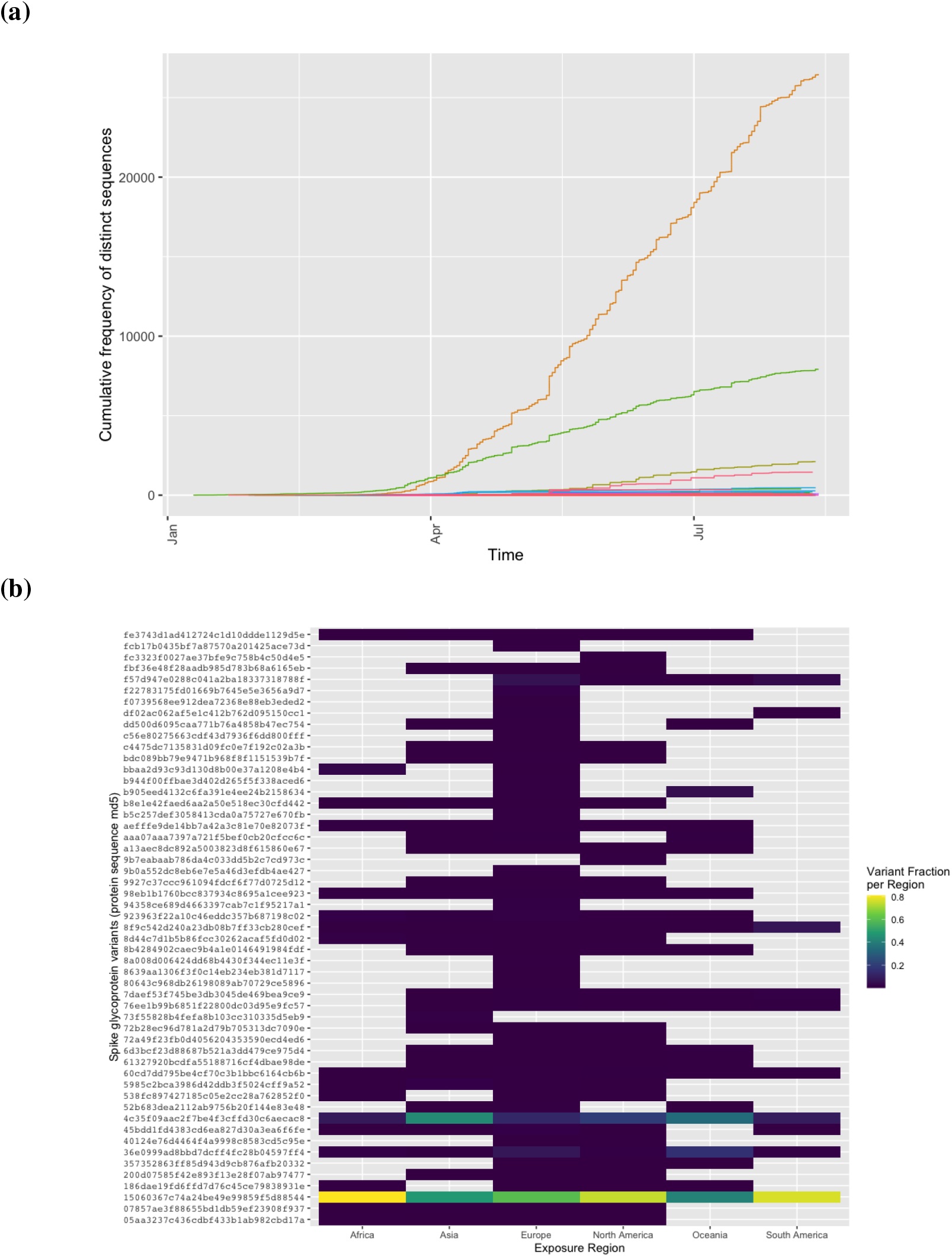
Spike glycoprotein variants observed in SARS-CoV-2 genomes over time and geography. Each line represents the cumulative frequency per variant (orange: D614G, green: UniProt ID P0DTC2, olive: P1140X, pink: S2 cleavage product) in **6a**. Low frequency S protein sequences (*<*5 observations) are removed from plotting for simplicity. In **6b**, the proportion of spike glycoprotein variants differ by exposure region. Proportion is calculated per variant to allow inter-region comparisons.

When S protein variants are stratified by region of exposure (Figure 6b), the ancestral variant (`uid key:15060367c74a24be49e99859f5d88544`) is the most predominant proportion globally across the corpus (0.42 – 0.80 of observed variants for a given exposure region) whereas the D614G variant (`uid key:4c35f09aac2f7be4f3cffd30c6aecac8`) is observed as a lower proportion per region (range 0.08 – 0.44) the next most prevalent variant. Both of these variants are observed across all exposure regions. Together, these results provide temporal and geographic insights about the dominant variants in the population of genomes that have been analyzed in this work. Additionally, our pipeline correctly identifies key D614G and N501Y variants that are been previously observed and experimentally validated (*28*) further indicating its accuracy.

## 3 Discussion

Since the start of the SARS-CoV-2 global pandemic, there have been immense efforts globally to sequence with near real-time efficiency the viral genomes observed in infected patients. In order to capitalize on this large and growing corpus of data, high throughput computational methods must be developed for rapid, high accuracy analysis to deliver the molecular targets that are actually under evaluation for drug development, vaccine specificity, and diagnostic testing. The method described here provides one such avenue to accomplish this goal. The protein and domain data generated as part of this work provides these molecular targets in an efficient manner with very high accuracy across the entire SARS-CoV-2 proteome and for all genomes analyzed in this corpus spanning multiple countries and lineages. Beyond this, our semi-supervised pipeline does not require the use of a single reference genome which better allows the detection of novel or mutating gene, protein, and respective domain sequences as they emerge. The method described here has been integrated with our Functional Genomics Platform and applied to hundreds of thousands of SARS-CoV-2 genomes. As the vaccination rates rise and the pandemic continues, this method can be used to rapidly monitor and track emerging protein variants to inform vaccine specificity and host protein binding affinity. Additionally as future work, further confirming the *in silico* predicted sequences using a structural model will allow for refinement of the protein sequences and key domains to expand our understanding of interaction with host proteins, antivirals, or diagnostics. Overall, the data generated as part of this work provides a comprehensive set of protein and domain variants observed globally and supports the research community as we aim to understand and control the SARS-CoV-2 pandemic.

## 4 Methods

We used a combination of state of the art tools and custom calibration tools to provide a semi-supervised genome annotation pipeline. We verified accuracy of, and applied this method to 66,905 SARS-CoV-2 genomes to identify the gene, protein, and functional domain sequences contained within each genome. This collection was analyzed for accuracy and quality with regard to current known references. Protein variants are characterized as a function of time since the pandemic emerged and from a geographic perspective.

### 4.1 Genome Data Retrieval and Quality Thresholds

SARS-CoV-2 genomes were retrieved from the Global Initiative for Sharing All Influenza Data (GISAID) (*19*) and NCBI GenBank (*18*) (retrieved August 18, 2020). A complete list of data sources, genome accessions, and acknowledgement of the submitting lab/author information where available is included in Supplemental File 1. An md5 hash was computed on each genome sequence (excluding headers) to track identical genome sequences. In preparation of genome annotation, two commonly used genome quality criteria and thresholds were assessed for their ability to yield a complete set of full length protein sequence annotations. Criteria A is defined as genome length *>* 29,000 bp (only IUPAC characters allowed, gaps permitted), % unknown bases (Ns) *<* 1, “high coverage” flag indicated by GISAID defined as *<* 0.05% mutation density only in CDS, and no unverified indels in relation to all other genomes in GISAID. Criteria B is defined as: number of unknown bases *<*= 15, number of degenerate bases *<*= 50, number of gaps *<*= 2, and mutation density *<* 0.25. For benchmarking genome quality criteria, all genomes were processed with our genome annotation pipeline and their resulting protein sequences were evaluated. Protein length distributions as a function of this genome criteria were compared using a two-sample Kolmogorov-Smirnov (ks.test function in base **R**).

### 4.2 Gene and Protein Annotation

Specific modifications to our previously described genome annotation pipeline (*20*) were made to process SARS-CoV-2 genomes and yield gene, protein, and domain sequences. In the sub-sections below we describe, in detail, the key modifications of Prokka v1.14.5 (*8*) for improved unsupervised annotation of SARS-CoV-2 genomes (Section 4.2.1) and the addition of custom-built supervised algorithms to improve identification of specific proteins that were unable to be detected using the base implementation (Section 4.2.2). The method has been Dockerized and is available for use at https://github.com/IBM/omxware-getting-started/tree/master/SARS-CoV-2_parser.

#### 4.2.1 Modifications to accommodate SARS-CoV-2 genome attributes and nascent state of reference data

To yield gene and protein names, Prokka (*8*) requires a reference protein database as a BLAST (*32*) index. We constructed this from the UniProt COVID-19 pre-release reference (ftp://ftp.uniprot.org/pub/databases/uniprot/pre_release/covid-19.dat). During the build phase for this index (Prokka script: `prokka-uniprot to fasta db`), the following modifications were made and applied during SARS-CoV-2 annotation:

1. Modify minimum evidence level required from transcript level (evidence=2) to predicted (evidence=4) when selecting reference proteins. This change allows proteins with evidence levels: at the protein level (evidence=1), at the transcript level (evidence=2), inferred from homology (evidence=3), or predicted (evidence=4) to be used when building references (but does not include protein uncertain, evidence=5). This is to better accommodate the nascent state of SAR-CoV-2 protein references.
2. Do not assign “hypothetical protein” to recommended full names that start with the following regular expression:

~~~
/^UPD\d|^Uncharacterized protein|^ORF|^Protein /
~~~

as some valid SARS-CoV-2 proteins contain these prefixes e.g. ORF3a protein and Uncharacterized protein 14.
3. Accept proteins without a recommended full name as long as the entry includes a full name provided by the submitter e.g. ORF10. Based on the above, the BLAST index was built using `prokka-uniprot_to_fasta_db` with the following command parameters: `--verbose --term Viruses –evidence`

`4.`The output of this command was then copied to the

`db/kingdom/Viruses` folder of the Prokka distribution to use as reference data.

Additionally, if a genome is less than 100,000 bp in length, by default Prokka will automatically switch to “metagenome mode” even in the absence of the metagenome flag opposed to persisting in “single genome mode.” To ensure single genomes of SARS-CoV-2 which are 29,000 bp were processed appropriately, we added an option named `mintotalbp` to the main `prokka` script which parameterized the minimum total base pairs required before this auto-switch could be activated. We set the default value for `mintotalbp` to 100 bp to avoid this inadvertent mode switch.

#### 4.2.2 Modifications to improve complete and accurate protein identification

Custom processing was developed for ORF9b, ORF10, Envelope small membrane protein, and Replicase polyprotein 1ab. To support additional identification of these sequences, all raw potential gene sequence coordinates and their scores were output from Prodigal using `-s` (opposed to only gene coordinates above Prodigal’s default threshold) during an intermediate step prior to Prokka with additional modifications as indicated below.

For the ORF9b, ORF10, and Envelope small membrane protein, an additional extraction process was completed by parsing that extended potential gene coordinate information. A length search was completed from these putative coordinates where accepted sequences must be within 10% of the reference protein sequence length and be the closest match to known reference sequences within those length bounds. Global alignments between the candidate protein sequence and known references (UniProt IDs: P0DTD2, A0A663DJA2, and P0DTC4 respectively) were completed using pairwise2 in BioPython with BLOSUM62 scoring matrix (*33*), gap open penalty = -2, and gap extend penalty = -1 to indicate sequence similarity between predicted and known reference sequences.

For Replicase polyprotein 1ab (pp1ab) important modifications were added beyond this to ensure identification of the full length sequence and to accommodate the naturally occurring -1 programmed ribosomal frameshift (*6*). We used the raw predicted gene coordinates from Prodigal to extract a candidate gene sequence from the originating genome. However, these candidate coordinates do not yield the full gene length or correct full length pp1ab sequence and were therefore modified as follows:

1. If Prodigal outputs two separate segments of the full gene sequence, we augmented and filled in the missing gap section from the originating genome based on the overall start and end coordinates to yield one contiguous gene sequence.
2. If Prodigal output only one truncated segment of the full pp1ab gene sequence, we shifted the starting index to ensure that the full length sequence achieved the expected entire 21,289 bp known to be part of the reference sequence (UniProt ID:P0DTD1).
3. In both cases, we verified that the gene sequence begins with an expected start codon (Methionine, ATG) and ends with a proper stop codon (TAA). When identifying the start codon, we verified the expected first three nucleotides were in the predicted sequence and shifted the start index to ensure this was the start position if that was not the case. Then, if the sequence did not include the start codon, we subtracted 1 from the start index until the correct start codon (ATG) was the first three nucleotides. The same procedure was used to ensure that the sequence ended with a proper stop codon, as we add 1 to the end index until TAA were the last three nucleotides.
4. Next, the slippery site as identified by Kelly, et al. (*6*) was identified in the gene sequence allowing for nucleotide degeneracy as indicated.
5. At the point of the slippery site, the preceding base was repeated and the remaining gene sequence was appended to yield the gene sequence which was then translated to yield the full length pp1ab protein sequence.

This method for generating complete Replicase polyprotein 1ab sequences was applied to all genomes. Of the 5,055 genomes below our quality control thresholds (inactive genomes), only 3,056 genomes were observed to contain a slippery site and therefore only those were able to be analyzed using this method.

### 4.3 Protein domain annotation

Unique protein sequences were processed with InterProScan v5.48-83 (*9*) to identify domain sequences and InterPro (IPR) codes as previously described (*20*). This version of InterProScan contains a number of InterPro, Gene Ontology and Pathway codes specific to the SARS-CoV-2 proteome and reference data. A full list of all available codes can be found at https://www.ebi.ac.uk/interpro/proteome/uniprot/UP000464024/.

### 4.4 Comparative analysis

To compare our method against other published viral genome annotation tools, VAPiD (v1.2 with Python3) was run on a set of 100 randomly selected SARS-CoV-2 genomes above quality control thresholds previously defined in Section 4.1 using the following parameters: reference (`--r`) NC 045512.2. Protein names and sequences were extracted from VAPiD output files using BioPython’s parser. Prokka version 1.14.5 (*8*) was run on this same set of genomes using default parameters with `--kingdom Viruses`. The resulting protein sequences from each tool were compared for set membership per genome, protein sequence truncations, and overall sequence similarity.

Protein annotations were evaluated against the SARS-CoV-2 proteome reference sequences indicated in ViralZone, SIB Swiss Institute of Bioinformatics (*22*) for complete protein set membership per genome, sequence length, and sequence similarity to known references indicated in NCBI UniProt (*21*). Set membership accuracy is the count of observations of a given protein for a set of genomes analyzed or in the case of domains, the set of domain sequences annotated for a given protein.

For domain accuracy comparative analysis, our predicted domains identified in spike glycoprotein (S protein) were analyzed for set membership completeness against the expected InterPro domain architecture for UniProt reference sequence P0DTC2 (https://www.ebi.ac.uk/interpro/protein/reviewed/P0DTC2/). Additionally, where predicted domain sequences were assigned an IPR code (8,146 unique domain sequences out of 9,120 total domain sequences), the predicted domain sequence was compared against the reference sequence to yield a percent identity. Reference domain sequences were extracted from the S protein amino acid sequence (UniProt:P0DTC2) based on domain start and stop sites indicated at the link above. Amino acid percent identity was calculated with considerations for insertions, deletions, or substitutions.

For genome to genome and protein variant comparisons (Sections 2.1 and 2.5), genome-associated metadata was retrieved from GISAID and processed for each analysis. For duplicated genome identification, an md5 of the genome sequence (excluding header) was completed as described in 4.1. Originating lab, date submitted, and host fields were used to further characterize candidate duplicate genome sequences. For protein variant analysis, the date submitted and exposure region fields are used to describe the time and geography of the observed variants.

### 4.5 Data Availability

The Functional Genomics Platform is available at

https://ibm.biz/functional-genomics. Access to the data generated from the method described herein is available through a developer toolkit (REST services, omxware Python SDK, and Docker container) or web interface, which can be accessed by requesting credentials at the link above. This includes the data described in this manuscript as well as the continual update of new identifications. Additionally pertaining to this manuscript, protein and domain sequence data are provided in Supplemental Files 2 and 4, respectively with identifier mappings described in Supplemental Files 3 and 5. All the GISAID data is available at www.gisaid.org.

## 5 Author Contributions

KLB, ES, and VM conceived of this work. KLB, ES, GN, and AA designed the experiments. KLB, ES, GN, AA, and HK generated and analyzed the data. KLB, ES, GN, and AA wrote the manuscript. SB, VM, JK analyzed the data and oversaw the experiments. All co-authors revised and approved the manuscript.

## Acknowledgments

We would like to acknowledge the immense collaborative effort from the IBM COVID-19 Task Force to help us open this platform to support research globally for the SARS-CoV-2 pandemic. Additionally, we gratefully acknowledge all the authors from the originating laboratories responsible for obtaining the specimens and the submitting laboratories where SARS-CoV-2 genetic sequence data were generated and shared via the GISAID Initiative, on which part of this research is based.

## Supplementary materials

- Supplementary File 1: SARS-CoV-2 Genome Acknowledgements (txt)
- Supplementary File 2: Non-redundant Protein Sequences (fasta.gz)
- Supplementary File 3: Genome and Protein Mapping Information (csv)
- Supplementary File 4: Spike Glycoprotein Domain Named Sequences (csv)
- Supplementary File 5: Protein and Domain Mapping Information (tsv)

## Footnotes

1 https://ibm.biz/functional-genomics

